## Supplementary Information for "Cell-TRACTR: A transformer-based model for end-to-end segmentation and tracking of cells"

- Supplementary Figures
- Supplementary Movie Captions
- Supplementary Text
- Supplementary Methods

#### Supplementary Figures

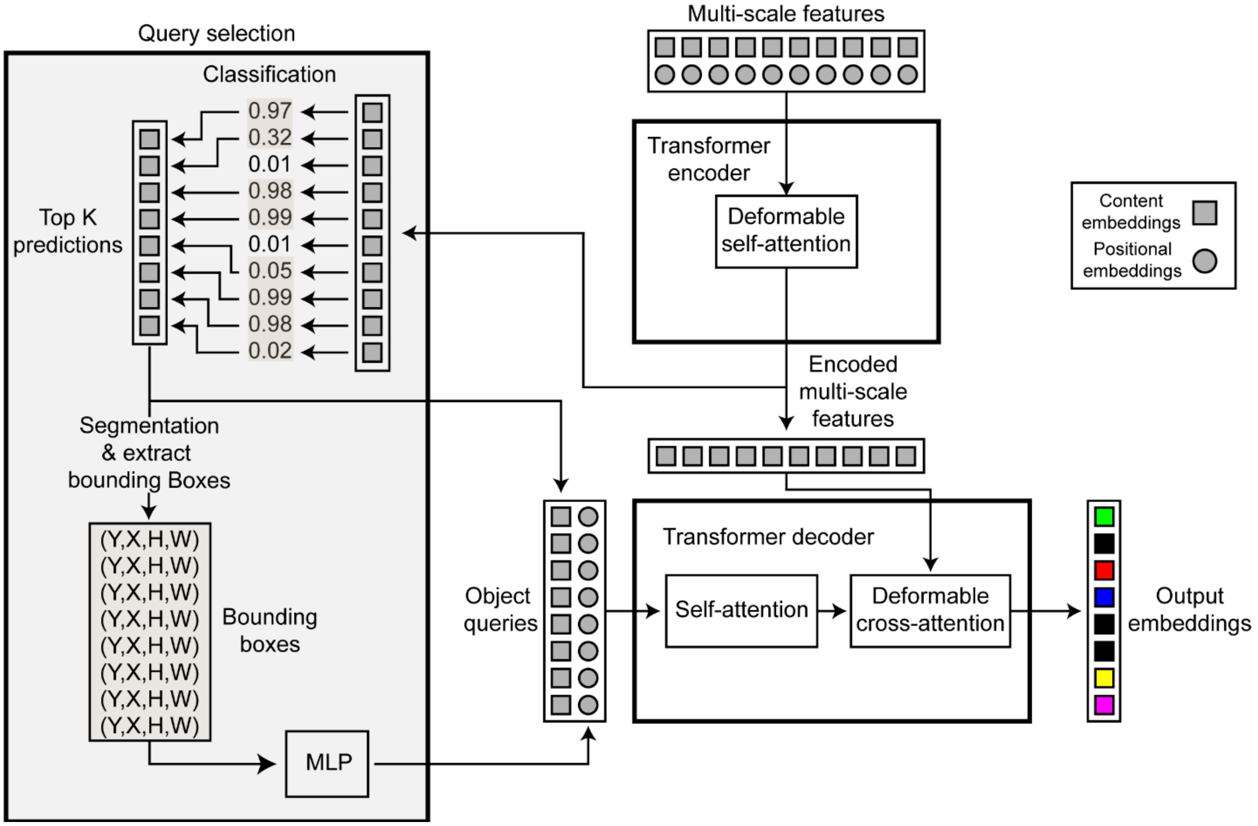


**Figure S1**. The encoder employs deformable self-attention to transform multi-scale features into encoded multi-scale features. Query selection is used to derive the object queries from these encoded multi-scale features. Each encoded image feature undergoes classification, and the top-K features are selected as the content embeddings for the object queries. Additionally, bounding boxes are extracted from the top-K segmentation masks and are subsequently converted into positional embeddings. Following this, the object queries perform self-attention and deformable cross-attention with the encoded multi-scale features to produce the output embeddings. MLP, multi-layer perceptron.

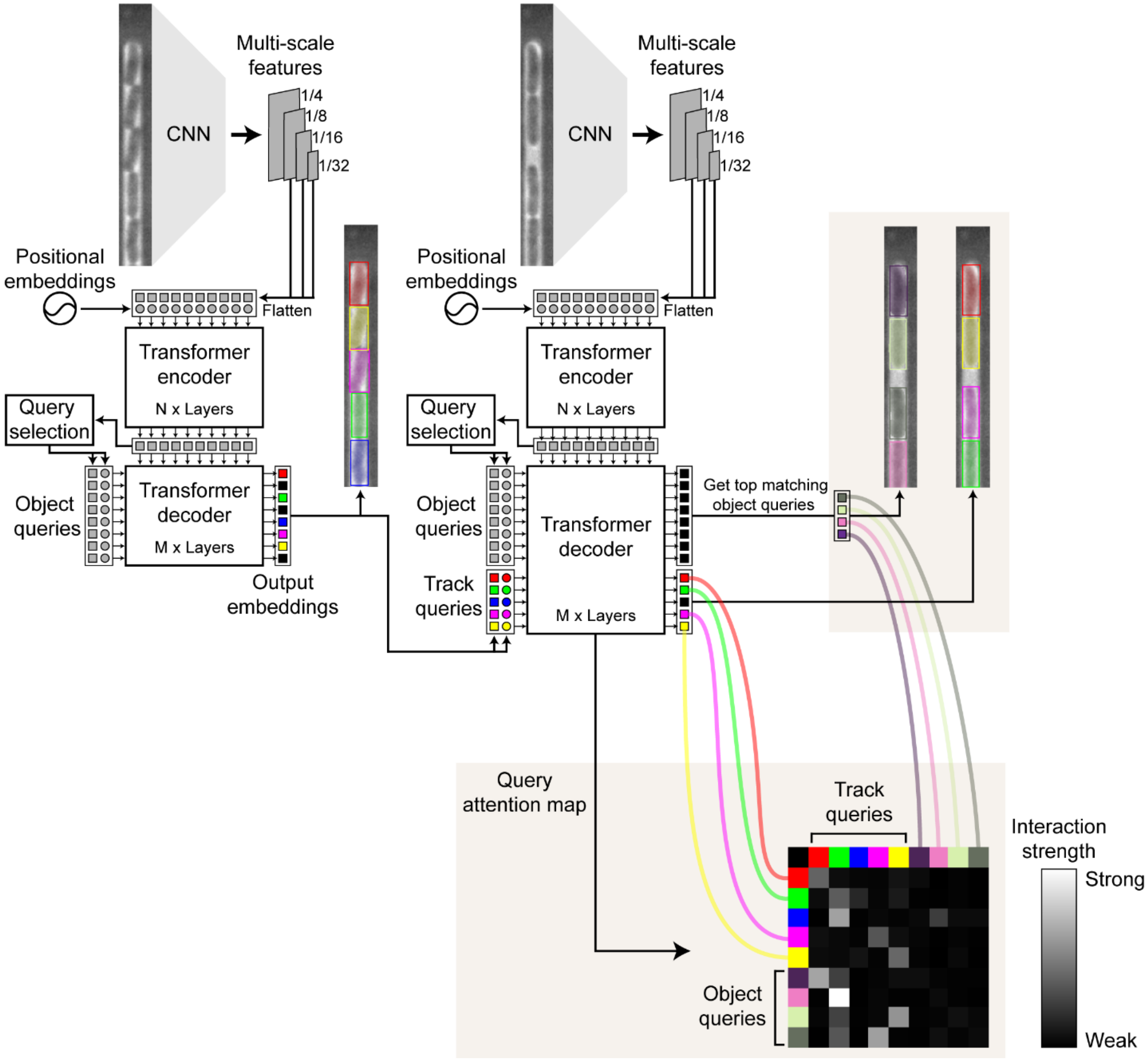


**Figure S2**. Example illustrating how object and track queries interact using an attention map processed by Cell-TRACTR across two subsequent frames. The schematic displays the actual output alongside the output the from top matching object queries. Extracted from the self-attention mechanism in the decoder, the attention map highlights the strength of interactions among the queries. Lighter colors indicate stronger interactions. The track queries tend to focus on themselves, while the object queries attend to the track query that corresponds most closely in location.

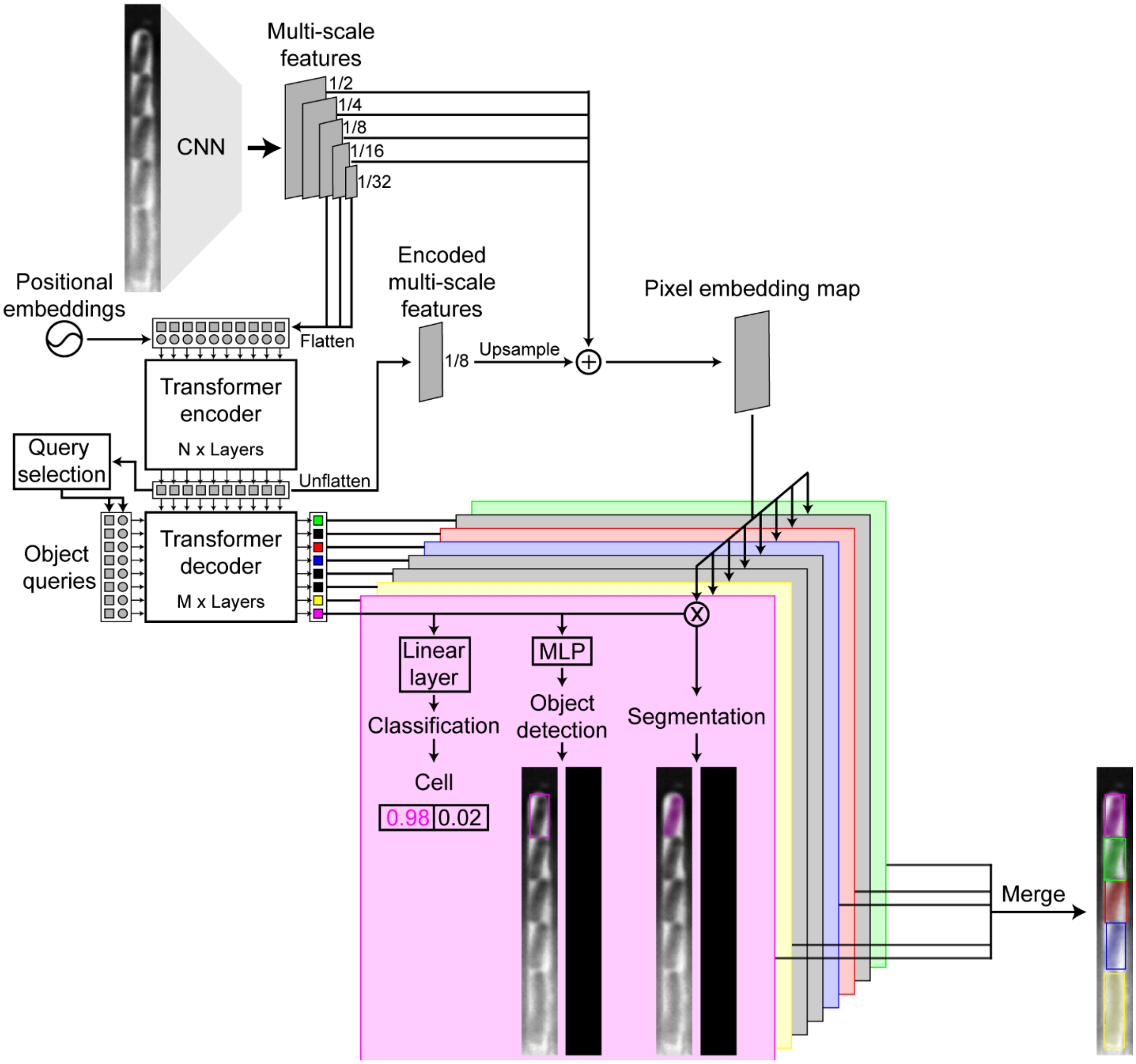


**Figure S3**. Depiction of how predictions are made from the output embeddings. The largest encoded multi-scale feature map is resized and combined with the original multi-scale features to form the pixel embedding map. The dot product of the output embeddings and the pixel embedding map produces the segmentation masks. A linear layer is used to predict the class label and a multi-layer perceptron (MLP) is used to generate the bounding box. For output embeddings classified as "no object", the bounding boxes and segmentation masks are ignored.

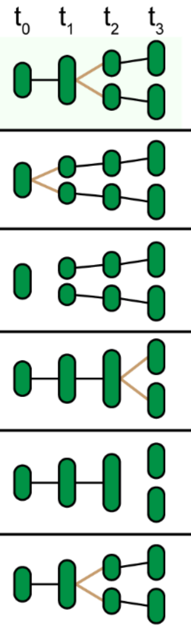


| | FP | FN | NS | EA | ED | EC | AOGM ↓ | TRA ↑ |
|---|---|---|---|---|---|---|---|---|
| Ground truth | | | | | | | | |
| Tracker A Early division | 2 | 1 | 0 | 3 | 4 | 0 | 17 | 0.75 |
| Tracker B Missing division | 2 | 1 | 0 | 3 | 2 | 0 | 15 | 0.78 |
| Tracker C Late division | 0 | 0 | 1 | 4 | 3 | 0 | 14 | 0.79 |
| Tracker D Missing division | 0 | 0 | 1 | 4 | 1 | 0 | 12 | 0.82 |
| Tracker E Correct | 0 | 0 | 0 | 0 | 0 | 0 | 0 | 1 |

**Figure S4**. Example showing how $OP_{CTB}$ can reward a tracker for predicting no division rather than predicting a division one frame too early or late. Acyclic oriented graph matching (AGOM) is the weighted sum of the operations needed to transform the tracker graph into the reference graph. A lower AGOM score is better and a higher TRA score is better.

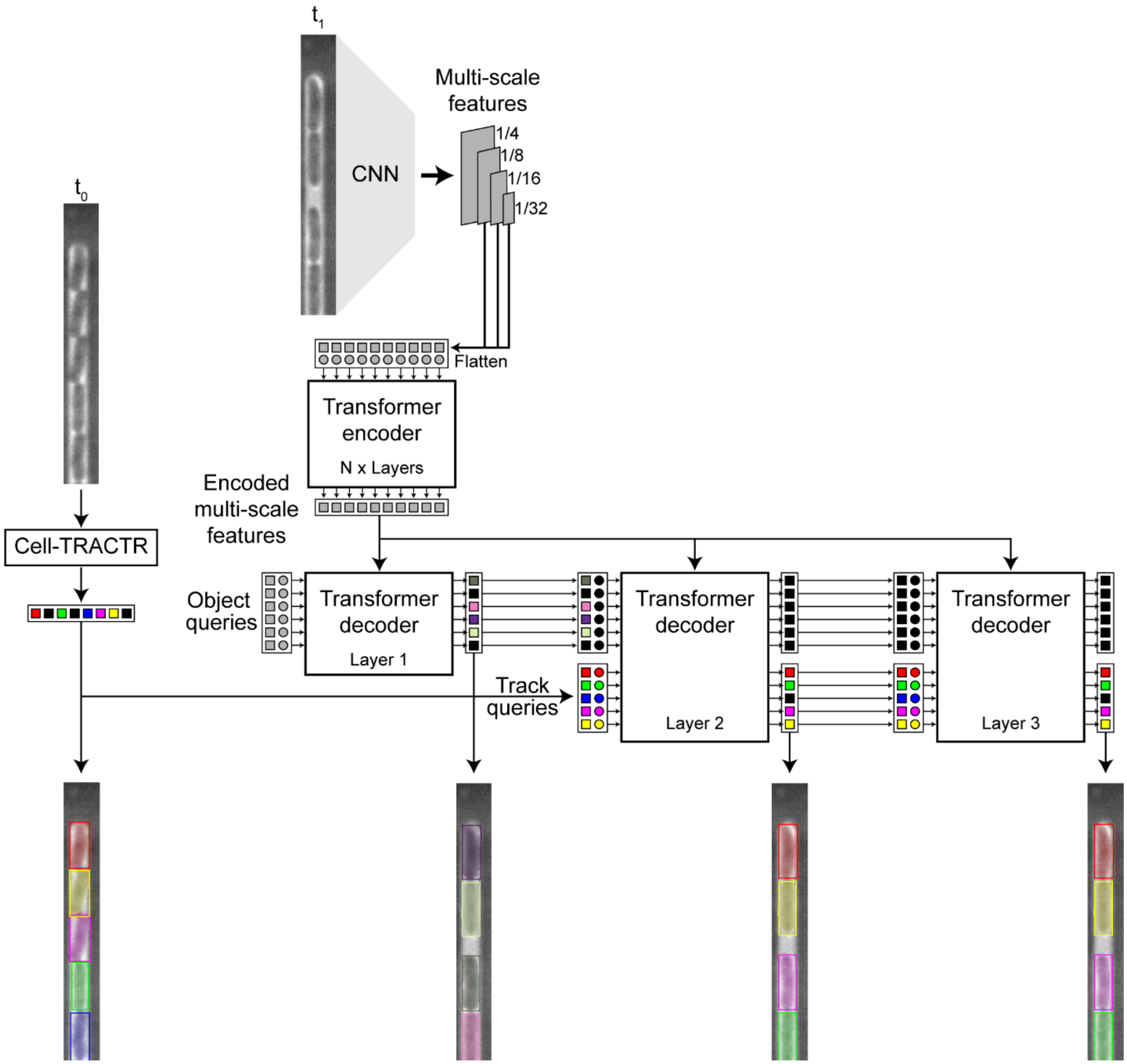


**Figure S5**. Cells are initially detected in frame $t_0$ and then tracked to frame $t_1$. Object queries are processed in the first layer of the decoder for object detection. In the subsequent decoder layers, track queries are concatenated with the object queries to perform tracking.

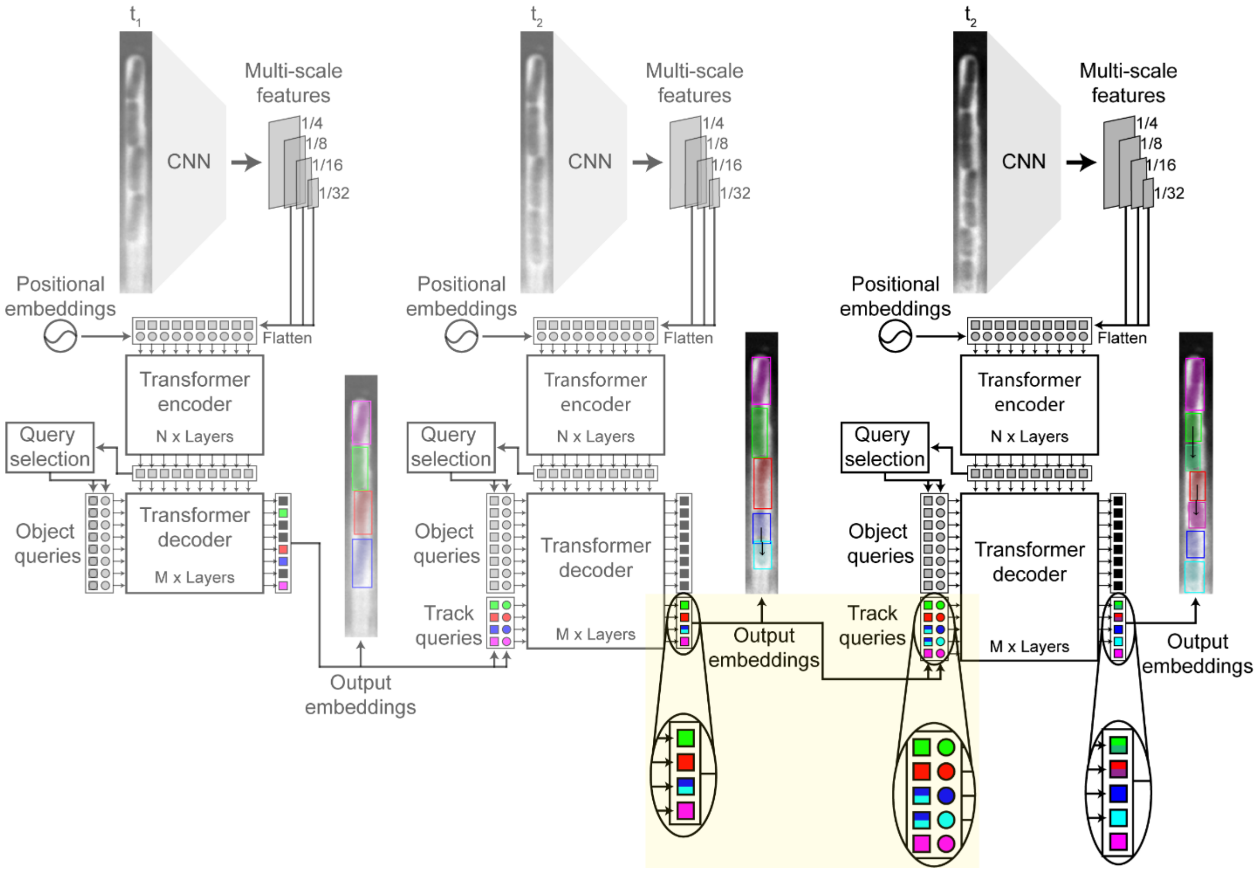


**Figure S6**. Schematic demonstrating how Cell-TRACTR tracks cells post division. The dark blue cell at time $t_1$ is predicted to divide at time $t_2$. The content embedding is duplicated for both divided cells. The positional embeddings are generated from each divided cell's segmentation mask. The content embeddings are shown as squares and the positional embeddings are circles.

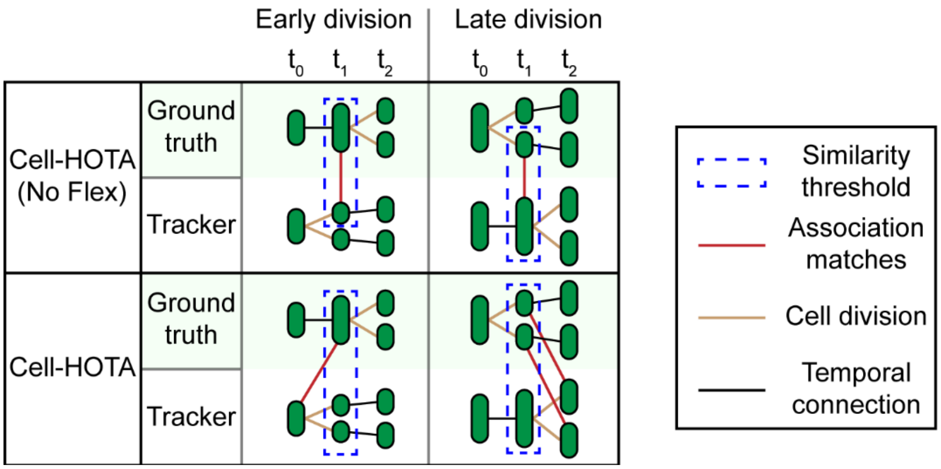


**Figure S7**. Schematic demonstrating how Cell-HOTA handles early and late divisions. For early divisions, the ground truth cell is originally matched with one of the tracker cells at time $t_1$. The red line indicates the matched cells, and the dotted blue box shows how the IOU is computed. To account for an early division, the IOU is computed between the ground truth cell in $t_1$ and both tracker cells in $t_1$. In addition, the temporal association for the ground truth cell in $t_1$ is altered to match the tracker cell in $t_0$. Similarly, for late divisions, the tracker cell is originally matched to one of the ground truth cells at time $t_1$. To account for a late division, the IOU is computed between the tracker cell in $t_1$ and both ground truth cells in $t_1$. The temporal associations are added between the ground truth cells in $t_1$ and the tracker cells in $t_2$. Note that for clarity this schematic only shows changes that are affected by the early or late division.

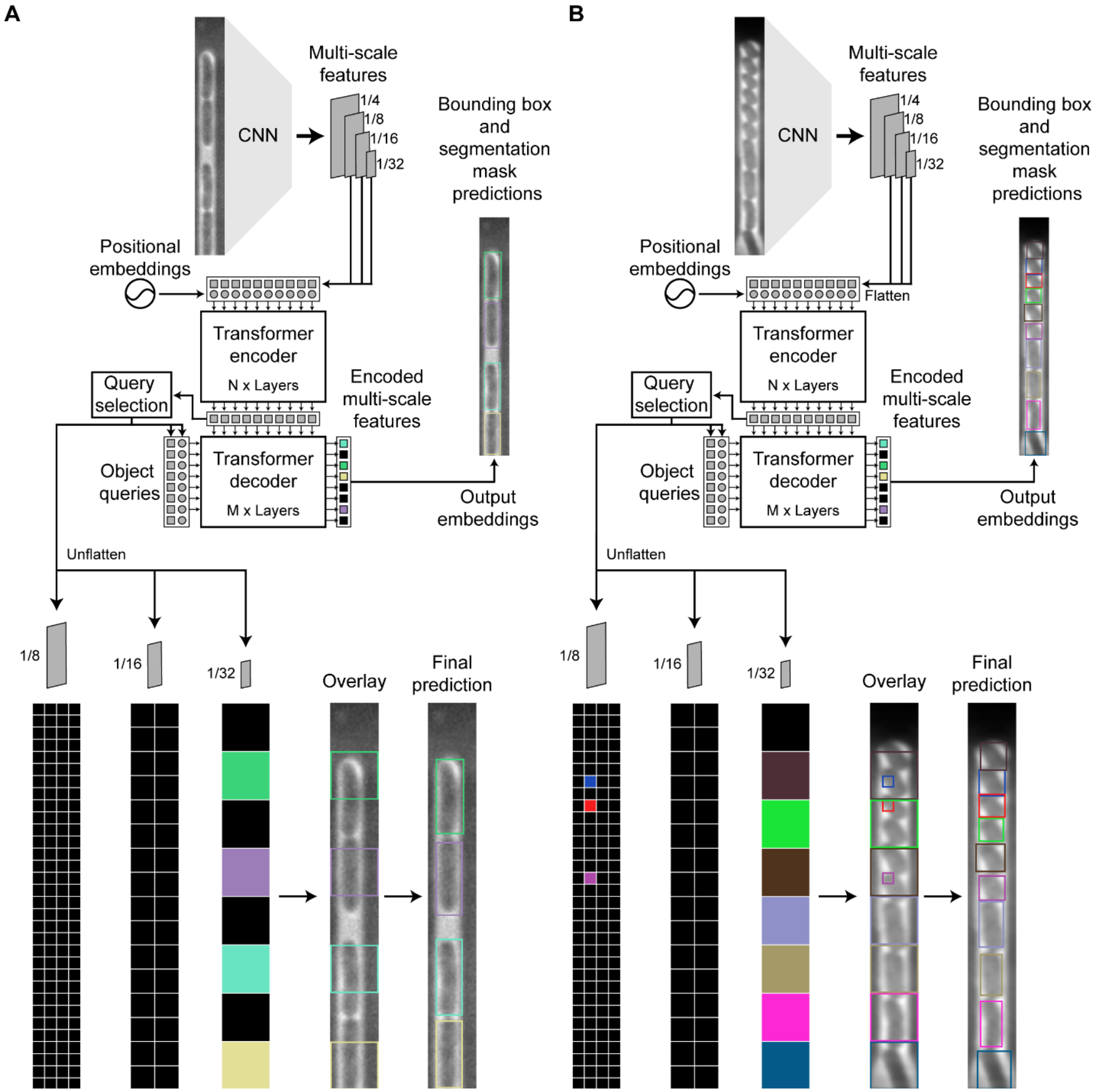


**Figure S8**. (A) Example illustrating the correlation between the final prediction of an object query and where the object query is derived from within the multi-scale features. Colored pixels represent multi-scale features that were classified as "cell" by the decoder. Black pixels represent multi-scale feature pixels that were classified as "no object" by the decoder. The colored pixels are overlaid onto the phase contrast image for visualization purposes. (B) A more complex example illustrating the correlation between the multi-scale features and object queries.

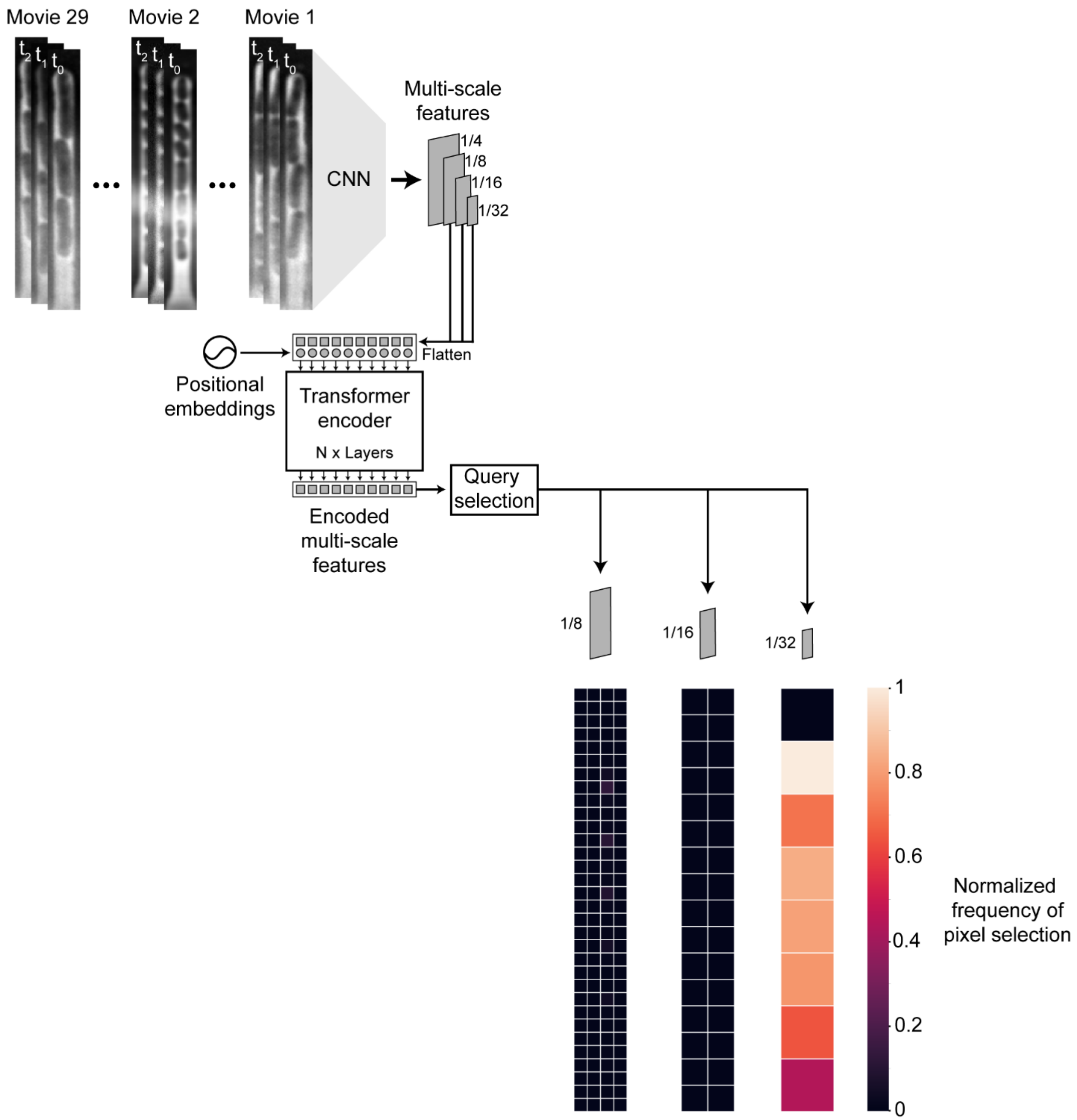


**Figure S9**. Heatmap illustrating which multi-scale feature pixels were classified as "cell" by query selection. This data was collected by processing the full test set containing 29 movies from the bacterial mother machine dataset with Cell-TRACTR. Note that for these movies, the model preferentially uses pixels from the lowest resolution multi-scale features. The black pixel at the top corresponds to the top of the mother machine image where there are not usually cells.

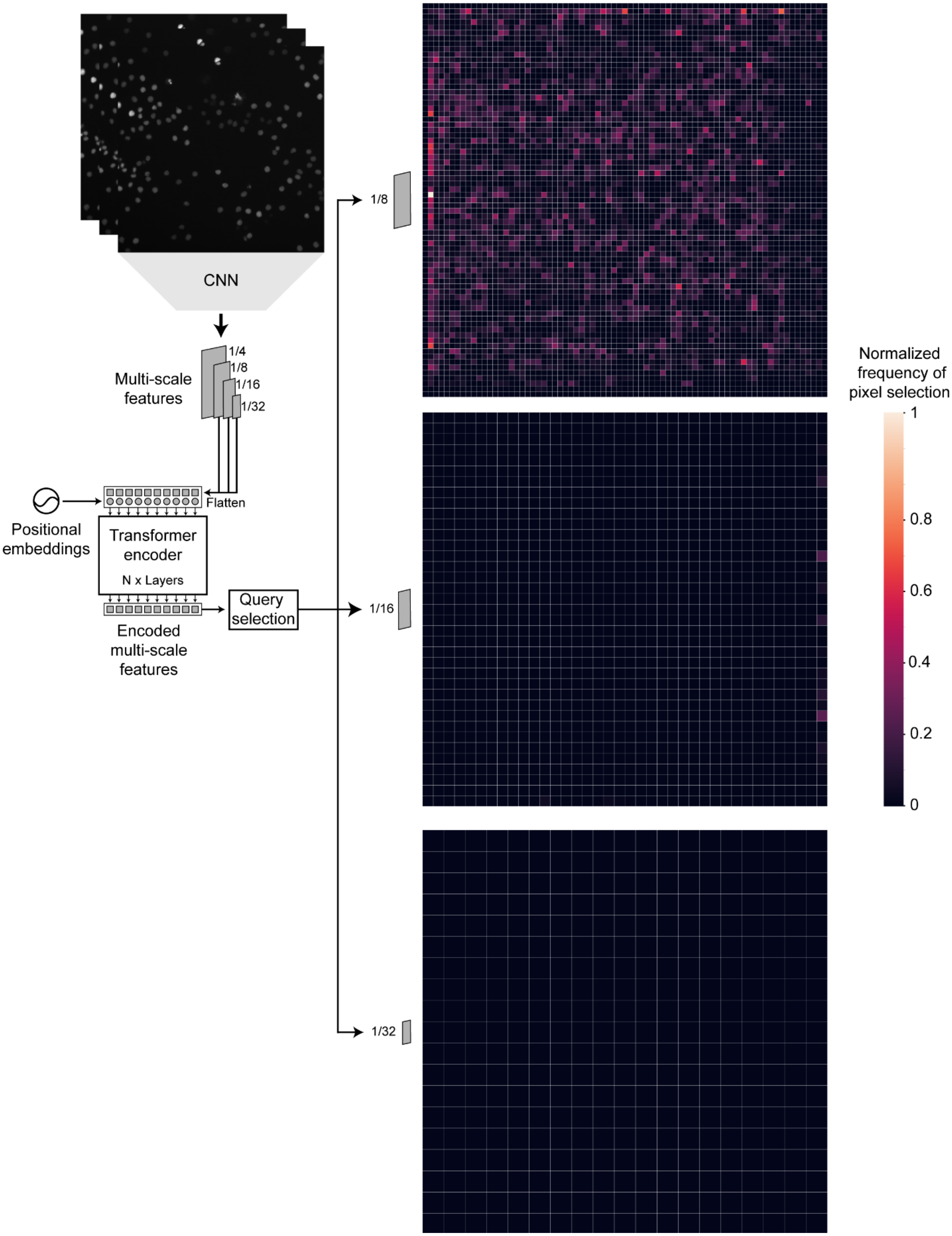

CNN
Multi-scale features
1/4
1/8
1/16
1/32
Flatten
Positional embeddings
Transformer encoder
N x Layers
Encoded multi-scale features
Query selection
1/8
1/16
1/32
Normalized frequency of pixel selection
1
0.8
0.6
0.4
0.2
0

**Figure S10**. Heatmap illustrating which multi-scale feature pixels were classified as “cell” by query selection. This data was collected by processing the full test set for the DeepCell dataset. Note that for these movies, Cell-TRACTR preferentially uses pixels from the highest resolution multi-scale features.

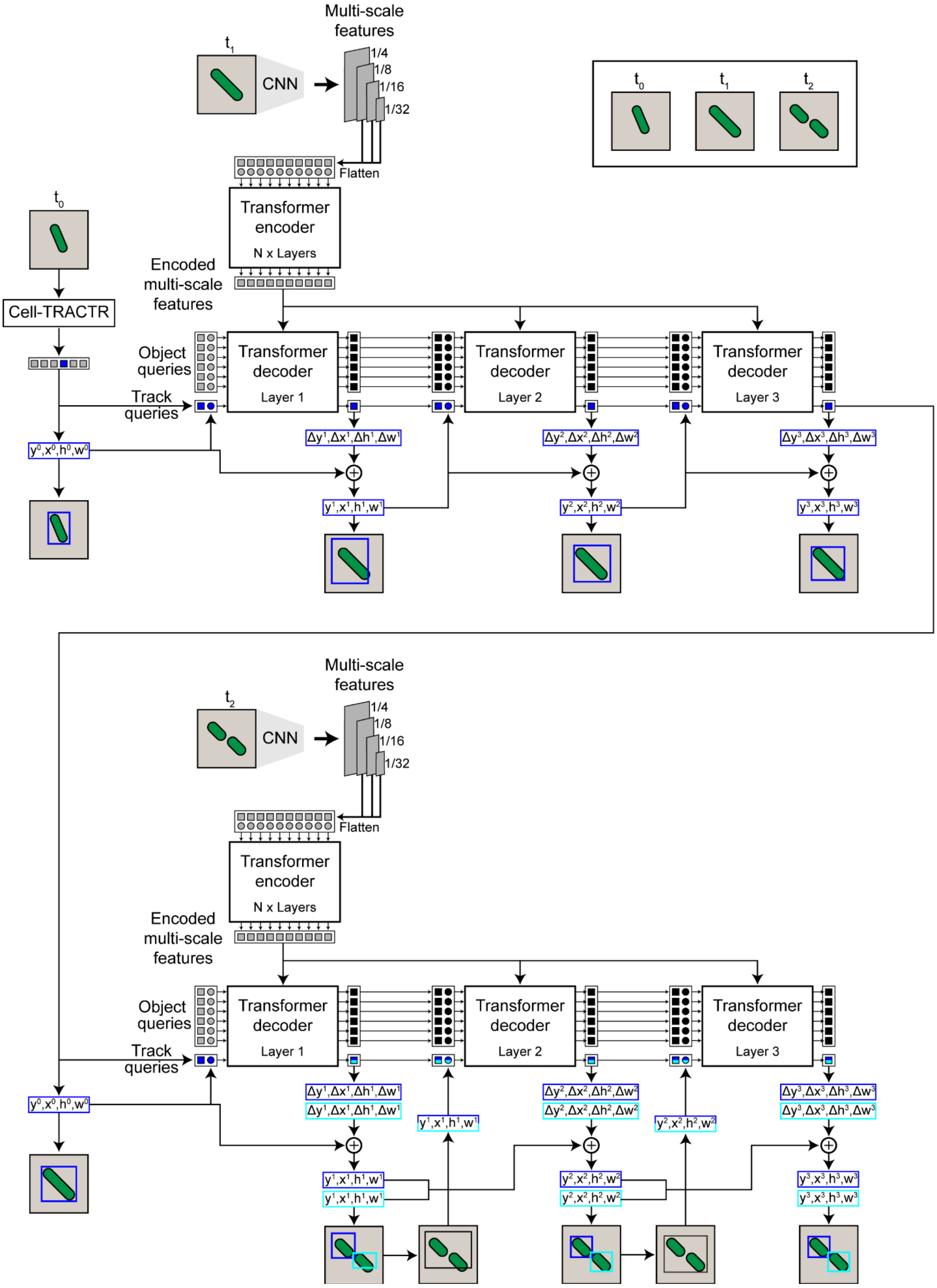

Multi-scale features
t1
CNN
1/4
1/8
1/16
1/32
Flatten
Transformer encoder
N x Layers
Encoded multi-scale features
t0
t1
t2
Cell-TRACTR
Object queries
Track queries
Transformer decoder
Layer 1
Layer 2
Layer 3
y0,x0,h0,w0
Δy1,Δx1,Δh1,Δw1
Δy2,Δx2,Δh2,Δw2
Δy3,Δx3,Δh3,Δw3
y1,x1,h1,w1
y2,x2,h2,w2
y3,x3,h3,w3
t2

**Figure S11**. Diagram illustrating how the iterative bounding box refinement method works with cell divisions. When tracking the cell from $t_0$ to $t_1$, the bounding box from $t_0$ serves as the initial bounding box that is iteratively refined three times into a bounding box that locates the cell in $t_1$. The second bounding box is ignored here since cell division is not predicted at time $t_1$. This method consists of predicting a relative offset with respect to the previous bounding box, represented by a change in cell location and size ($\Delta y$, $\Delta x$, $\Delta h$, $\Delta w$). This offset is added to the previous bounding box to get the new prediction. After the first layer in the decoder, the predicted relative offset is added to the initial bounding box in $t_1$. This new bounding box serves as the reference point for the next layer. The reference point indicates where the query should focus within the encoded multi-scale features. In the two subsequent layers, the predicted relative offset is added to the previous bounding box to get the final bounding box location. At time $t_2$, the model predicts a cell division resulting in both relative offsets being utilized. After the first layer in the decoder, the predicted relative offsets are added to the same initial bounding box in $t_2$. In the two subsequent layers, the two predicted relative offsets are added to their respective previous bounding box. We generate a new bounding box around both divided cells that serves as the new reference point. This only occurs when both class labels for a track query are greater than 0.5.

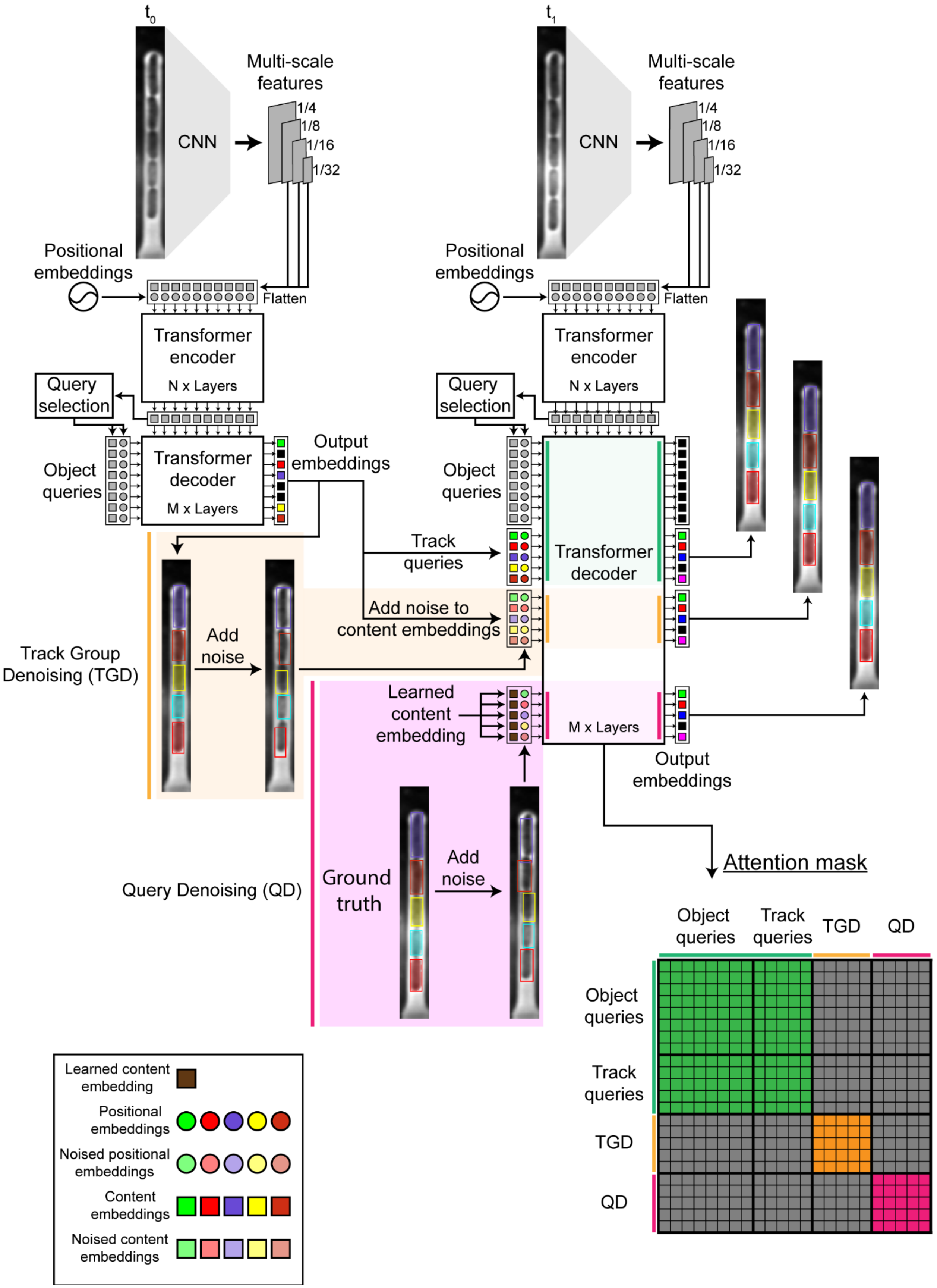

t0
CNN
Multi-scale features
1/4
1/8
1/16
1/32
Positional embeddings
Flatten
Transformer encoder
N x Layers
Query selection
Object queries
Transformer decoder
M x Layers
Output embeddings
t1
Track queries
Add noise to content embeddings
Track Group Denoising (TGD)
Add noise
Learned content embedding
Query Denoising (QD)
Ground truth
Output embeddings
Attention mask
TGD
QD
Learned content embedding
Positional embeddings
Noised positional embeddings
Content embeddings
Noised content embeddings

**Figure S12**. Cell-TRACTR detects cells in the first frame at time $t_0$. Object queries predicted to be a "cell" are converted to track queries. For Track Group Denoising (TGD), random noise is added to the track queries to get noised track queries. For Query Denoising (QD), random noise is added to the ground truth bounding boxes to generate noised track queries. All queries are processed together by the decoder. Attention masks are used to prevent information leakage between the object and track queries, the noised track queries generated from TGD, and the noised track queries generated from QD. The gray boxes indicate information being blocked between sets of queries while colored boxes indicate the flow of information.

#### Supplementary Movie Captions

**Movie S1**. Time-lapse microscopy movie from the bacterial mother machine test set processed by Cell-TRACTR. Different colors represent unique cells being tracked. Black arrows indicate cell division. The temporal resolution of the movie is one frame every five minutes. Both the bounding boxes and segmentation masks are displayed for visualization purposes. However, during inference, only the segmentation masks are used.

**Movie S2**. Time-lapse microscopy movie from the mammalian DeepCell test set processed by Cell-TRACTR. Different colors represent unique cells being tracked. Black arrows indicate cell division. The temporal resolution of the movie is one frame every five minutes. Both the bounding boxes and segmentation masks are displayed for visualization purposes. However, during inference, only the segmentation masks are used.

**Movie S3**. Time-lapse microscopy movie from the bacterial mother machine test set processed by Cell-TRACTR and evaluated with the Cell Tracking Challenge (CTC) $OP_{CTB}$ metrics. From left to right, the first image shows the raw images, the second image shows the ground truths, and the third image shows the predictions. Distinct colors represent unique cells being tracked over time. The right six images show the operations used to transform the predicted graph into the reference graph provided by the ground truth. The operations include *delete vertex* (FP), *add vertex* (FN), *split vertex* (NS), *add edge* (EA), *delete edge* (ED), and *alter the edge semantics* (EC). When one of these operations is applied, the cell is shown in red.

### Supplementary Text

***Cell Tracking Challenge metric ($OP_{CTB}$) and limitations for frequently-dividing cells***

The Cell Tracking Challenge metric for quantifying overall performance of the cell tracking benchmark, $OP_{CTB}$ (Equation S1), incorporates measures of both tracking accuracy (TRA) and segmentation accuracy (SEG).

$$OP_{CTB} = 0.5 * (TRA + SEG) \ (S1)$$

Acyclic oriented graphs are employed to compute the TRA score. The spatial locations of cells are denoted by vertices while the edges signify the temporal associations (1). For each series of time-lapse images, an acyclic oriented graph is generated based on the output from a tracking algorithm. These graphs are then compared to a reference graph which contains the information from the ground truth. The acyclic oriented graph matching (AOGM) measure is a weighted sum of the number of operations needed to transform the predicted graph into the reference graph (Equation S2). These operations include *split vertex* (NS), *add vertex* (FN), *delete vertex* (FP), *delete edge* (ED), *add edge* (EA), *alter the edge semantics* (EC), where we follow the notation used by the Cell Tracking Challenge. The AOGM is a weighted sum of the number of the operations required to match the two graphs.

$$AOGM = w_{NS}NS + w_{FN}FN + w_{FP}FP + w_{ED}ED + w_{EA}EA + w_{EC}EC \ (S2)$$

where $w_{NS} = 5$, $w_{FN} = 10$, $w_{FP} = 1$, $w_{ED} = 1$, $w_{EA} = 1.5$, and $w_{EC} = 1$. $AOGM_0$ is the cost needed to generate the reference graph from scratch. AOGM is normalized to $AOGM_0$ and subtracted from 1 to get the TRA score (Equation S3). To ensure the score remains positive, the AGOM score cannot be greater than the $AOGM_0$ score. If tracking is perfect, resulting in AGOM = 0, then the tracking score TRA = 1.

$$TRA = 1 - \frac{\min(AOGM, AOGM_0)}{AOGM_0} \ (S3)$$

As in the main text, we define a 'tracker cell' as a cell predicted by the tracking algorithm and a 'ground truth cell' as a cell defined in the ground truth. In the case of perfect tracking, each tracker cell would be uniquely affiliated with one ground truth cell, however errors in tracking cause deviations from this. In the $OP_{CTB}$ metric, a tracker cell needs to overlap with at least half the area of a ground truth cell to be considered a match. A tracker cell without a match is a false positive and a ground truth cell without a match is a false negative. A tracker cell can match with multiple ground truth cells, but a ground truth cell can only match with at most one tracker cell. The *split vertex* operation is used when a tracker cell matches with multiple ground truth cells whereas the *add vertex* and *delete vertex* operations are used for false negatives and false positives, respectively. The operations *add edge* and *delete edge* are used to add and remove temporal connections between cells. Lasty, the *alter the edge semantics* operation modifies existing temporal connections.

The Cell Tracking Challenge uses the SEG score to measure localization accuracy. The Jaccard Similarity index is used to calculate the intersection over union (IOU) between each ground truth cell (GT) and matched tracker cell (TrCell) (Equation S4).

$$J(\text{TrCell}, GT) = \frac{|GT \cap \text{TrCell}|}{|GT \cup \text{TrCell}|} \quad (S4)$$
$$where \quad |GT \cap \text{TrCell}| > 0.5 * \ |GT|$$

The SEG score is the Jaccard Similarity index averaged over all the ground truth cells (Equation S5).

$$SEG = \frac{1}{N_{GT}} \sum_{1}^{N_{GT}} J(\text{TrCell}, GT) \quad (S5)$$

There are some drawbacks to using $OP_{CTB}$, especially in cases where cell division events are frequent, and the timing of division is not precisely known. In some cases, removing division links, thus breaking cell lineages that should remain intact, can increase the TRA score despite being a worse tracking prediction. As a reminder, a higher TRA score is better than a lower TRA score. For example, a division event predicted a frame early or late relative to the ground truth can result in a lower TRA score than the equivalent case with division entirely removed (Fig S4). This decrease in score occurs because the predicted division creates edges that need to be removed to recreate the reference graph. When examining Cell-TRACTR's performance on a movie from the test set, 5 of 5 *split vertex* operations and 20 out of 21 *add edges* operations occurred when a cell division was predicted late by one frame (Movie S3). Thus, a significant portion of the error used to calculate the TRA score comes from early and late divisions which is undesirable for evaluating tracking performance since these are somewhat arbitrary distinctions because it is challenging to generate ground truths that pinpoint the exact frame a cell divides. As a result, test sets used to evaluate the cell tracking algorithms are bound to contain ambiguous cell division events. The TRA score will measure the algorithm's ability to exactly match imperfect ground truths instead of measuring the algorithm's ability to track cell divisions at a broader level.

***$OP_{CTB}$ versus HOTA***

HOTA offers several potential benefits over the $OP_{CTB}$ score. With $OP_{CTB}$, the TRA and SEG scores are completely independent of each other. The TRA score serves as the detection and tracking accuracy whereas the SEG score provides the localization accuracy. The TRA score is reported at one specific similarity threshold $\alpha$. Therefore, it is hard to gauge how the tracking algorithm will perform with a stricter or looser similarity threshold. Although the SEG score provides information about the localization accuracy, it is difficult to relate this to the TRA score. In contrast, the HOTA score integrates the localization accuracy into DetA and AssA by evaluating them at multiple similarity thresholds.

Another issue with the $OP_{CTB}$ is that the TRA score lacks interpretability. The TRA score uses detection operations (*add vertex*, *remove vertex*, *split vertex*) that are comprehensible, however the tracking operations (*add edge*, *remove edge*, *alter the edge semantics*) do not clearly translate into understandable errors. In addition, the *split vertex* operation is used to correct late divisions however there is no inverse relationship. Plus, it is difficult to specifically evaluate division accuracy, which is critical for cell tracking. Conversely, Cell-HOTA breaks down into DetA, AssA, and DivA which facilitates straightforward evaluation of the algorithm's capacity to track cells. For example, if a small percentage of cells in a movie are detected but they are all tracked correctly, then DetA would be low and AssA would be high. However, if most cells in a movie are detected but they are tracked incorrectly, then DetA would be high and AssA would be low. Interpretability of these scores is crucial for figuring out how to improve an algorithm.

Lastly, the $OP_{CTB}$ is weighted heavily towards localization accuracy, as the score is computed as the average of the SEG and TRA scores, where SEG represents the localization accuracy and TRA represents the detection and association accuracy. Ideally, a tracking metric should balance the detection, association, and localization accuracy. The HOTA score effectively addresses this by weighing the DetA and AssA equally and incorporating localization accuracy into both of those metrics.

***Directly comparing Cell-HOTA and $OP_{CTB}$ results***

In order to directly compare Cell-HOTA with TRA from $OP_{CTB}$, we need to look at the Cell-$HOTA_{0.5}$ score. Cell-$HOTA_{0.5}$ measures the overall detection and tracking accuracy at $\alpha = 0.5$. Therefore, a tracker cell and ground truth cell only match if the IOU is greater than 0.5. Similarly, the TRA score measures the overall detection and tracking accuracy where a tracker cell needs to overlap with at least half the area of the ground truth cell to be considered a match. When assessing model performance on the mammalian DeepCell dataset, Cell-TRACTR achieves the highest Cell-$HOTA_{0.5}$ score by a large margin yet had the lowest TRA score. Even when ignoring $DivA_{0.5}$, which would most closely resemble the TRA score since it puts little weight on cell division, Cell-TRACTR still attains the highest $DetA_{0.5}$ and $AssA_{0.5}$. This discrepancy is due to how the TRA score accounts for false positives and false negatives. The Cell-HOTA metric gives equal weight to false negatives and false positives whereas the TRA score gives ten times the weight to false negatives compared to false positives. The weight configuration for the AGOM measure used to generate the TRA score was designed to reflect the effort needed to correct an error manually (1). When analyzing the TRA score on the mammalian DeepCell dataset, EmbedTrack accrued 4080 false positives and 150 false negatives whereas Cell-TRACTR had 1262 false positives and 918 false negatives. Although Cell-TRACTR had significantly fewer false positives and false negatives combined, the high weight towards false negatives had a more significant impact on the TRA score.

#### Supplementary Methods

***Query selection***

To give object queries more context, query selection generates region proposals from the encoder output (2) (Fig S1). Each pixel in the multi-scale features acts as a potential region proposal, predicting a bounding box, segmentation mask, and class label. The top-K scoring region proposals are selected to serve as object queries for the decoder. For region proposals that are classified as "cell", there is a strong correlation between the pixel location of the regional proposal within the multi-scale features and the predicted location of the cell in the final layer of the decoder (Fig S8). In general, we found that Cell-TRACTR preferentially used the low-resolution multi-scale features for the bacterial mother machine dataset (Fig S9) and high-resolution multi-scale features for the mammalian DeepCell dataset (Fig S10). Following Mask-DINO (3), segmentation masks are used to generate the positional embeddings for the object queries. To minimize computational load, backpropagation is performed only for the top-K scoring region proposals.

***Adding noise to bounding boxes during training***

Random noise ($\Delta x$, $\Delta y$) is generated by selecting two numbers from a uniform distribution ranging between -1 to 1. We use noise coefficients $\lambda_1$ and $\lambda_2$, with default values of 0.2 and 0.1 respectively, for center shifting and box scaling respectively. These noise coefficients scale the amount of random noise added to the boxes. Center shifting affects the (x, y) coordinates of the box, whereas box scaling affects the height and width of the box.

***Matching algorithm***

During training, we use the Hungarian algorithm to match each ground truth to a prediction (4). For object detection, each ground truth cell is optimally matched to a predicted cell using this approach. This optimization is based on the class label, bounding box, and segmentation mask similarities. When cells are being tracked, the predicted cells are automatically matched with their respective ground truths. The remaining ground truth cells which are not tracked, like cells entering the field of view, are matched to a predicted cell derived from an object query. For cell division, a set of two predicted cells need to match with a set of two ground truth cells. However, this could be problematic as the daughter cells are matched as a set, not as individual cells. The order of the daughter cells should match the order of the ground truth cells. Predicted daughters could be matched to the ground truth daughters of a reversed order. To address the ambiguity of the order in cell division, we use the Hungarian algorithm to pick the correct order of ground truth cells.

***Reference points for cell divisions***

Most DETR-like models that track objects do not deal with dividing objects (5–9). Therefore, in classical DETR-like models, each query will only predict one class label, bounding box, and segmentation mask. Below, we describe the approach that is used in DETR-like models which we also employ in Cell-TRACTR; in the following paragraph we discuss the extensions we made to handle cell division that are specific to Cell-TRACTR. In the decoder, the bounding boxes are converted into positional embeddings to assist in self-attention and cross-attention. However, the raw bounding boxes are also used as reference points for cross-attention. Queries attend to a small set of sampling points around the reference point (2). Instead of each query attending to the whole feature map, it will attend to a specific area on the feature map. Deformable attention reduces time to convergence and computational cost. Since deformable attention was designed for image processing, it is not used during self-attention. For each layer in the decoder, iterative bounding box refinement is performed where the decoder refines the bounding box prediction. These bounding boxes are then used to generate the reference points and positional embeddings needed for the next layer.

In Cell-TRACTR, there are two predictions made per query (Fig S11). For object queries, only the first prediction is utilized. However, track queries may predict a cell division. For track queries that predict a cell division, the combined bounding box between the two divided cells is used to generate the reference point and positional embedding. Like the original method, the decoder refines the bounding box prediction for each of the divided cells separately. The combined bounding box is used to generate the reference points and positional embeddings needed for the next layer.

***Alleviating the conflict between object and track queries***

While simultaneously detecting and tracking cells makes DETR-based models powerful, it also has drawbacks. Various studies have highlighted that conflict between object and track queries can lead to an overall decrease in detection accuracy (6–9). MOTRv2 (6) improves on MOTR (10) by utilizing the YOLOX (11) object detector to provide the positional embeddings for the object queries so the model can focus on association. MOTRv3 (7), which builds on top of MOTR (10), employs a Release-Fetch Supervision strategy that utilizes one-to-one matching between all queries and all objects in the first five layers and then one-to-one matching only between the object queries and newborn objects in the last layer. MeMOTR (8) simply uses the first decoder layer for object detection only and uses the subsequent decoder layers for joint detection and tracking. Similar to MOTRv3, CoMOT (9) implements Coopetition Label Assignment where the first five decoder layers perform an additional one-to-one matching with just the object queries and all objects (newborn and tracked) and then a standard last layer. Following these advances, we adopt the first decoder layer as a detection only layer like MeMOTR (Fig S5) and add an extra one-to-one matching between object queries and all objects in the subsequent layers except for the last layer, similar to the approach used in CoMOT (Fig S2).

***Track Group Denoising (TGD) and Query Denoising (QD)***

For faster convergence during training, we added denoised track group queries and denoised track queries as proposed in MotrV3 (7) and MotrV2 (6). Both training methods consist of the model learning to track noisy queries. Since the model simultaneously processes the original track queries alongside both the noised track group queries and noised track queries, it is important to regulate information flow across these training techniques. For example, the original track queries could cheat by leveraging information about a denoised track query. To prevent this, we employ attention masks to selectively block the attention mechanism from accessing certain parts of the input data (12,13) (Fig S12). Specifically, the masks are added to the attention scores before the softmax operation, setting the scores for the blocked parts to a very large negative value. This ensures that these parts have a negligible effect after the softmax, effectively isolating the attention to focus only on the relevant, unmasked portions of the data.

For TGD, random noised is added to the bounding boxes predicted from the previous frame and converted into positional embeddings (Fig S12). Following Ref. (6), we use noise coefficients of $\lambda_1 = 0.2$ and $\lambda_2 = 0.1$. Random noise, drawn from separate normal distributions with a mean of 0 and standard deviation of 0.1, is added to the content embeddings. TGD teaches the model to generalize and become less dependent on precise positional and content information. Unlike MotrV3, to reduce computational load we use only one track group, whereas they use multiple track groups. This TGD method, inspired by Group-DETR (14), has been shown to increase overall tracking performance (7).

For QD, all ground truth bounding boxes in the previous frame are collected. A small amount of noised is added to each bounding box (Fig S12). The noise coefficients, $\lambda_1 = 0.2$ and $\lambda_2 = 0.1$, are the same as used in TGD. The noised bounding boxes are then converted into positional embeddings. The same learned content embedding is used for all noised track queries. Due to the lack of semantic information, the model is forced to denoise the noised track queries solely based on the positional information. This method, which was introduced in DN-DETR (13), helps the model generalize and speed up training.

TGD and QD are very similar in implementation but differ in terms of targeting training efficiency and tracking performance. For TGD, positional embeddings originate from the predicted bounding boxes while for QD, the positional embeddings originate from the ground truth boxes. In addition, for TGD, the predicted content embeddings are utilized, whereas for QD, all noised track queries use the same learned content embedding. At a high level, TGD primarily improves tracking performance while QD primarily speeds up training. TGD and QD are only used during training. As a result, they do not add any computational load during inference.
